# The estrogenic pathway modulates non-breeding female aggression in a teleost fish

**DOI:** 10.1101/2020.02.12.946327

**Authors:** Lucía Zubizarreta, Ana C. Silva, Laura Quintana

## Abstract

Aggressive behaviors are widespread among animals and are critical in the competition for resources. The physiological mechanisms underlying aggression have mostly been examined in breeding males, in which gonadal androgens, acting in part through their aromatization to estrogens, have a key role. There are two alternative models that contribute to further understanding hormonal mechanisms underlying aggression: aggression displayed in the non-breeding season, when gonadal steroids are low, and female aggression. In this study we approach, for the first time, the modulatory role of estrogens and androgens upon non-breeding aggression in a wild female teleost fish. We characterized female aggression in the weakly electric fish *Gymnotus omarorum* and carried out acute treatments 1 h prior to agonistic encounters with either an aromatase inhibitor or an antagonist of androgen receptors. Aromatase inhibition caused a strong distortion of aggressive behavior whereas anti-androgen treatment had no effect on behavior. Territorial non-breeding aggression in female *G. omarorum* is robust and depended on rapid estrogen actions to maintain high levels of aggression, and ultimately reach conflict resolution from which dominant/subordinate status emerged. Our results taken together with our own reports in males and the contributions from non-breeding aggression in bird and mammal models, suggest a conserved strategy involving fast-acting estrogens in the control of this behavior across species. In addition, further analysis of female non-breeding aggression may shed light on potential sexual differences in the fine tuning of social behaviors.

**Highlights:**

- Female *Gymnotus omarorum* displayed robust territorial aggression in lab settings.
- Acute treatment with aromatase inhibitor lowered aggression levels.
- Aromatase inhibition increased first attack latency and decreased conflict resolution.
- Acute treatment with anti-androgens showed no effects.
- This is the first report of estrogens underlying teleost non-breeding female aggression.

## Introduction

Aggressive behaviors are widespread among animals, and they are key in the competition for resources such as food, shelter, and mating opportunities. Males are usually more aggressive than females, however robust female aggression is highly prevalent in many species, and not only in the context of maternal aggression. In particular, territorial aggression has been shown to occur in female fish, reptiles, birds, rodents, and non-human primates [1].

The physiological mechanisms underlying aggression have been mostly examined in breeding males, in which the involvement of gonadal androgens has been widely established [2]. In the last 30 years the understanding of the modulation of this complex behavior by sexual hormones has greatly advanced: estrogens have been recognized as additional modulators of aggression and both androgens and estrogens have been shown to have slow and rapid behavioral effects, reflecting genomic and nongenomic mechanisms ([3]; revised in [4]).

Researchers have incorporated two models which offer valuable opportunities to further understand the physiology of aggression: the very understudied female aggression, which is modulated by androgens and estrogens albeit frequently in ways distinct from males, and species in which aggression occurs uncoupled from the breeding season [5–13]. Female aggression has been shown to be promoted by testosterone and at least part of this effect is through its aromatization to estrogens [11]. Although estrogens may increase aggression in some species [9,10,14] their effects may differ and brain estrogen receptor subtypes have been shown to mediate opposing effects upon aggressive behavior [15,16]. Some species display aggression uncoupled from the breeding season, when their gonads are regressed and their circulating levels of gonadal androgens are reduced ([17]; revised in [13]). In the non-breeding season, estrogens have a forefront role in the regulation of aggression, mostly through rapid nongenomic mechanisms. Estradiol treatment has been shown to rapidly promote male non-breeding aggression in mammals and birds [18–23], and acute inhibition of aromatase, the enzyme which converts androgens into estrogens, decreases aggression levels [24]. In turn, aggressive interactions between males during the non-breeding season can produce changes of estradiol levels in specific brain areas [25]. In both males and females, non-breeding aggression is linked to modulations of brain estrogen receptors mediating rapid effects [18,26].

Androgens and estrogens can be synthesized locally in the brain either from extragonadal precursors [13,27–29] or *de novo* from cholesterol [30,31]. The study of non-breeding aggression of males and females has opened new avenues of understanding the complexity of the control of aggression and its overall modulation during natural seasonal cycles.

The weakly electric fish *Gymnotus omarorum* is a seasonal breeder, which displays year-long active territorial defense maintaining territories both in the natural habitat [32] and in the lab [33]. The non-breeding territorial aggression of wild *G. omarorum* is robust, elicited in neutral arenas and triggered by the presence of a conspecific [34]. It displays strikingly aggressive encounters, in both intra and intersexual dyads, and males and females show no differences in contest outcome, temporal dynamics of the agonistic encounter, levels of aggression, nor submissive signaling [34,35]. Male aggression has been rigorously characterized; contest resolution is biased by body size and once dominant/subordinate status is established; the dominant fish displays a long-lasting exclusion of the subordinate fish from its territory [33]. Male non-breeding aggression is independent of gonadal hormones, as it occurs robustly in fish that have been gonadectomized a month prior. In addition, intact fish show circulating androgen (11-ketotestosterone) levels unaffected by aggressive encounters. However, aggression is dependent on rapid hormone effects, as the acute inhibition of aromatase distorts contest dynamics and outcome: aggression levels are reduced, and outcome becomes unpredictable [36]. Is fast estrogenic modulation a general strategy underlying non-breeding aggression in this species, independently of sex? This study approaches wild female non-breeding aggression and its control by estrogens and androgens to address this issue.

## Methods

### Animals

We used wild adult females of *Gymnotus omarorum* (Richer-de-Forges et al., 2009) (body-length 15 - 26 cm and body weight 9 – 60 g) captured from the field and housed for 4 to 5 weeks in our facilities before experiments. All experiments were carried out during the non-breeding season (June to August) [37]. Fish were collected from Laguna del Sauce (34°51’S, 55°07’W), Maldonado, Uruguay using an electrical detector as previously described [37]. Animals were housed in individual mesh compartments (40×40×60 cm) within large outdoor tanks (500 L). These outdoor tanks house aquatic plants brought from the field and were subjected to conditions with natural photoperiod (from LD 10:14 to LD 11:13), temperature (10.41 ± 3.48 °C), and rainfall. To conserve conditions similar to the natural habitat, conductivity was maintained under 200 μS/cm [37]. Each fish had a shelter in its compartment and was fed *ad libitum* with *Tubifex tubifex*. All experiments were performed according to the regulations for the use of animals in research and the experimental protocol was approved by the institutional Ethical Committee of Instituto de Investigaciones Biológicas Clemente Estable (Resolution CEUA IIBCE 004/05/2016).

### Behavioral set up

We observed the agonistic behavior of *G. omarorum* in dyadic female-female encounters and tested the effect of aromatase inhibition or androgen receptor antagonism during the non-breeding season. The dyads (7-20% body weight difference between contenders) belonged to one of the following experimental groups: control dyads (n = 8), fadrozole-treated dyads (n = 10), or cyproterone acetate-treated dyads (n = 7). We performed the characterization of female agonistic behavior in control dyads. The evaluation of agonistic behavior included engagement in conflict, contest outcome, dynamics, aggression, and submission levels, and these parameters were used in the comparison to fadrozole and cyproterone acetate dyads. All experimental groups were composed of fish spanning the same size range and each fish was used only once. Dyads were placed in a behavioral setup (as described in [38]) that allowed simultaneous video and electric recordings, control of photoperiod, water temperature, conductivity, and pH. The setup consisted of 4 experimental tanks (55 × 40 × 25 cm) divided in half by a removable glass gate. Due to the nocturnal habits of this species, all experiments were performed at night, in darkness, with infrared LED illumination (Kingbright L- 53F3BT; 940 nm) located above the tanks. Experiments were recorded with an infrared-sensitive video camera (SONY CCDIris, Montevideo, Uruguay) through the glass bottom of the tank. The electric signals of freely moving fish were detected by two pairs of fixed copper wire electrodes connected to two high-input impedance (1 MΩ) amplifiers (FLA-01; Cygnus Technologies Inc., Delaware Water Gap, PA, USA). Images and electric signals were captured by a video card (Pinnacle Systems, PCTV-HD pro stick) and stored in the computer for further analysis. We used a neutral arena protocol with a plain arena (without food or shelter) and simultaneously placed each contender in one of the equally sized compartments 2 h prior to the experiment thus providing equal resources (territory and residency) to each individual [34]. Pharmacological manipulations were performed 1 hour before gate removal (see below). The gate was raised 10 min after sunset, and fish were separated 10 min after conflict resolution. Dyadic contests that did not reach an establishment of dominance/subordination after 20 minutes of interaction were interrupted and labeled as “dyads with engagement without resolution”. Dyadic interactions in which there was no engagement during 20 minutes after gate removal were interrupted and considered “dyads with no engagement”.

### Pharmacological manipulations

To analyze the rapid modulation of estrogens and androgens we used acute treatment with different inhibitors. Cyproterone acetate (CA) has been previously reported to effectively block AR in teleost fish [39,40]. Nevertheless, since it had never been used in *G omarorum*, we confirmed the innocuity of its vehicle and the effectiveness of the inhibitor in this species. We performed an experiment blocking a well-known androgenic-dependent trait previously described in non-breeding adults [41]. The electric organ discharge (EOD) of *G. omarorum* has a multiphasic waveform with four successive components (V1 to V4) [42]. A 15-day treatment with testosterone implants specifically increases the amplitude of the negative component V4 which is quantified by the index V4 amplitude/V3 amplitude (AV4/AV3). The reports on the effects of supraphysiological testosterone on EOD waveform were based on mixed groups (non-breeding males and females [41]). We subcutaneously implanted 22 animals with testosterone silastic pellets (100 ug/gbw). A stock solution of CA (Sigma, C3412), 2 μg/μl was prepared in mineral oil (Drogueria Industrial Uruguaya), and stored at 4°C. Testosterone implanted animals were divided into a control group (n = 12) which received a daily IP injection of mineral oil for 15 days, and the treatment group (n = 10) which received a daily IP injection of CA (20 μg/gbw) for 15 days. EOD waveform was recorded following [43]. The testosterone + mineral oil group increased AV4/AV3 amplitude index when comparing day 15 to day 0, as expected (paired t-test, p = 0.006, n = 12, Fig. 1A), whereas the group treated with testosterone + cyproterone acetate showed no significant increase in its index (paired t-test, p = 0.8, n = 10, Fig. 1B). This result demonstrates cyproterone acetate effectively blocks androgenic actions in *G. omarorum*, as shown in other teleost species. To verify the innocuity of mineral oil in the species we compared 6 females injected with PBS to 6 injected with mineral oil (in equal volumes). Fish were recorded individually in the behavioral setup, locomotion was quantified in a 2 minute time window 1 hour after injections. Percentage of time in movement was compared between oil and PBS females, showing no significant difference (Mann-Whitney U test, p = 0.2, n_OIL_ = 6, n_PBS_= 6). Basal EOD rate was calculated for each fish 1 hour after injections (see below), and there was no significant difference between oil and PBS females (Mann-Whitney U test, p = 0.57, n_OIL_ = 6, n_PBS_= 6).

**Figure 1.**
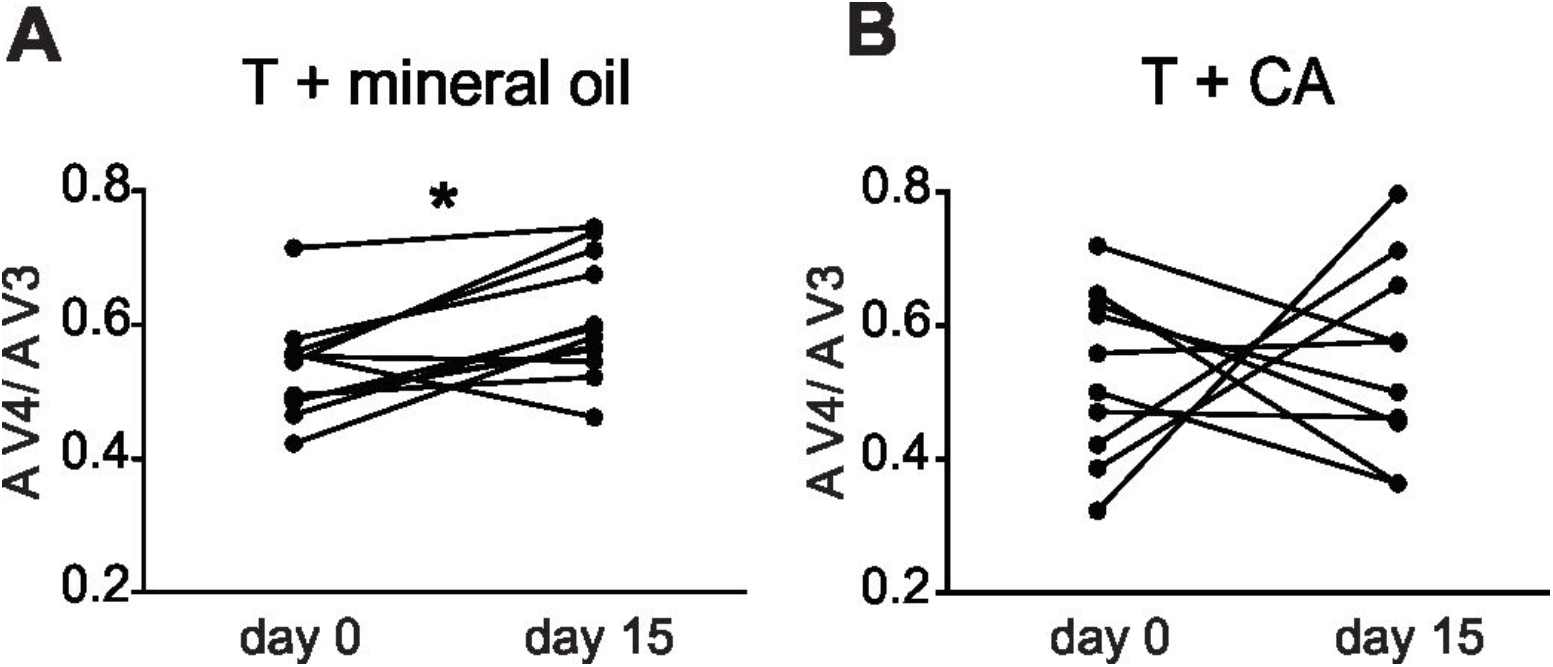
Test of effectiveness of cyproterone acetate (CA) in *Gymnotus omarorum*. **A**. Animals implanted with testosterone (T) and subjected to a daily IP injection of mineral oil (n=12) for 15 days changed their EOD waveform as expected [41], increasing the amplitude of the V4 component in comparison to day 1, shown as a significant increase of the index V4 amplitude / V3 amplitude (AV4/AV3) (paired t-test, p = 0.006, n = 12). **B.** Animals implanted with T and subjected to a daily IP injection of CA for 15 days (n=10) showed no significant differences in V4 amplitude (paired t-test, p = 0.8, n = 10).

To test the effect of acutely manipulating the androgenic pathway on agonistic behavior, we administered cyproterone acetate to female-female dyads before subjecting them to the neutral arena protocol. One hour before the agonistic encounter, we injected cyproterone acetate (10 μg/gbw, IP) to both individuals, behavioral experiments were performed as described above. To ensure an effective blocking we used a higher dosis than previously reported for other teleost species [39].

To assess the effect of modulating the estrogenic pathway we used the aromatase inhibitor fadrozole (FAD, Sigma F3806). Fadrozole has been previously reported to effectively block aromatase activity in teleost fish and other vertebrates, including this species [36,44–46]. Before the agonistic encounter, we diluted stock fadrozole (10 μg/μl) in PBS, and IP injected 20 μg/gbw to both individuals 1 hour prior to gate removal. Behavioral experiments were performed as described above. Control experiments in female-female dyads were carried out injecting PBS (in equivalent volume) to both contenders, one hour before the agonistic encounter. Individuals were sexed either by surgical observation 1 month prior, which has been shown has no effect upon agonistic behavior in comparison with intact dyads (FAD, control groups following [36]) or by gonadal inspection in euthanized animals after behavioral tests (CA group, euthanization by eugenol solution 8 mg l^−1^).

### Data processing

Locomotor and electric displays were analyzed by a researcher blind to the experimental groups and treatments. Following [34], we identified the three phases of agonistic encounters: (1) evaluation phase: from time 0 (gate removal) to the occurrence of the first attack; (2) contest phase: from the occurrence of the first attack to conflict resolution (resolution time); and (3) post-resolution phase: 10 min after conflict resolution. Conflict resolution was defined as the moment we observed the third consecutive retreat of one fish without retaliation. This criterion unambiguously defined subordinate status; fish fulfilling this requirement were never observed to change their status in the following 10 min of interaction. To calculate attack rate, we divided the number of attacks (bites, nips, nudges) [47] by contest duration time in seconds. We identified previously described transient submissive electric signals: offs (EOD interruptions), chirps (abrupt increases in EOD rate) [34] and electrical submission (stable, post-resolution EOD rate rank) [48]. We calculated off and chirp rate (for contest and post-resolution phases together) by dividing the number of offs and chirps produced in both phases by the duration in sec. To calculate electrical submission, we determined mean EOD rates in dominants and subordinates during pre-contest (before gate removal) and post-resolution in 10 - 60 s recordings from both phases, using the software Clampfit (Axon, 10.0.0.61). To quantify the difference in EOD rate between contenders, we calculated the subordinate / dominant EOD rate index (S/D rate index). Index values below 1 indicate that the dominant EOD rate is higher than the subordinate EOD rate.

### Statistics

To analyze the effect of cyproterone acetate on EOD waveform we used a paired t-test and compared A V4/ A V3 index in the same individual at day 0 and day 15 of treatment. As behavioral data did not fit a gaussian distribution, they were analyzed with non-parametric tests: Wilcoxon Matched-Pairs test (paired variables in the same fish or the same dyad comparing dominant and subordinate), Mann-Whitney U test (independent variables using sets of data from different fish). For this reason, results are expressed as median ± interquartile range throughout. Chi square test 2×2 Fisher was used to compare the proportion of dyads that achieved contest resolution in control and FAD groups.

## Results

### Female-female non-breeding territorial aggression

Female-female dyads of *G. omarorum* displayed robust agonistic behavior in the neutral arena protocol (Fig. 2). All dyads engaged in agonistic interactions, and all ended in the establishment of stable dominance/subordination relationships. The larger fish became dominant in 6 out of 8 contests. Agonistic encounters exhibited characteristic phases previously described for the species: (1) a short evaluation phase (first attack latency = 34.8 ± 8.8 s, n = 8); (2) a contest phase (contest duration, 273 ± 85.6 s, n = 8), and (3) a post-resolution phase (Fig. 2A). The contest phase was characterized by overt aggressive displays, higher in dominants compared to subordinates (attack rate Wilcoxon Matched-Pairs test, p = 0.008, n = 8, Fig 2B). In addition, dominant and subordinate attacks were strongly correlated during contests (R^2^ = 0.8, p = 0.003, n = 8, data not shown). During contest and post-resolution phase subordinates emitted electric signals of submission (off rate 0.02 ± 0.005, n = 8; and chirp rate 0.025 ± 0.01, n = 8, data not shown). After resolution, EOD rate rank was established, and the acquired status of dominants and subordinates did not reverse (Fig. 2C). In the pre-contest phase contenders did not differ in their basal EOD rates (median S/D rate index = 1.01) whereas after resolution dominants’ EOD rates were higher than their counterpart subordinates (median S/D rate index = 0.6; pre-contest vs post-resolution S/D rate index: Wilcoxon Matched-Pairs test, p = 0.016, n = 7, Fig. 2C).

**Figure 2.**
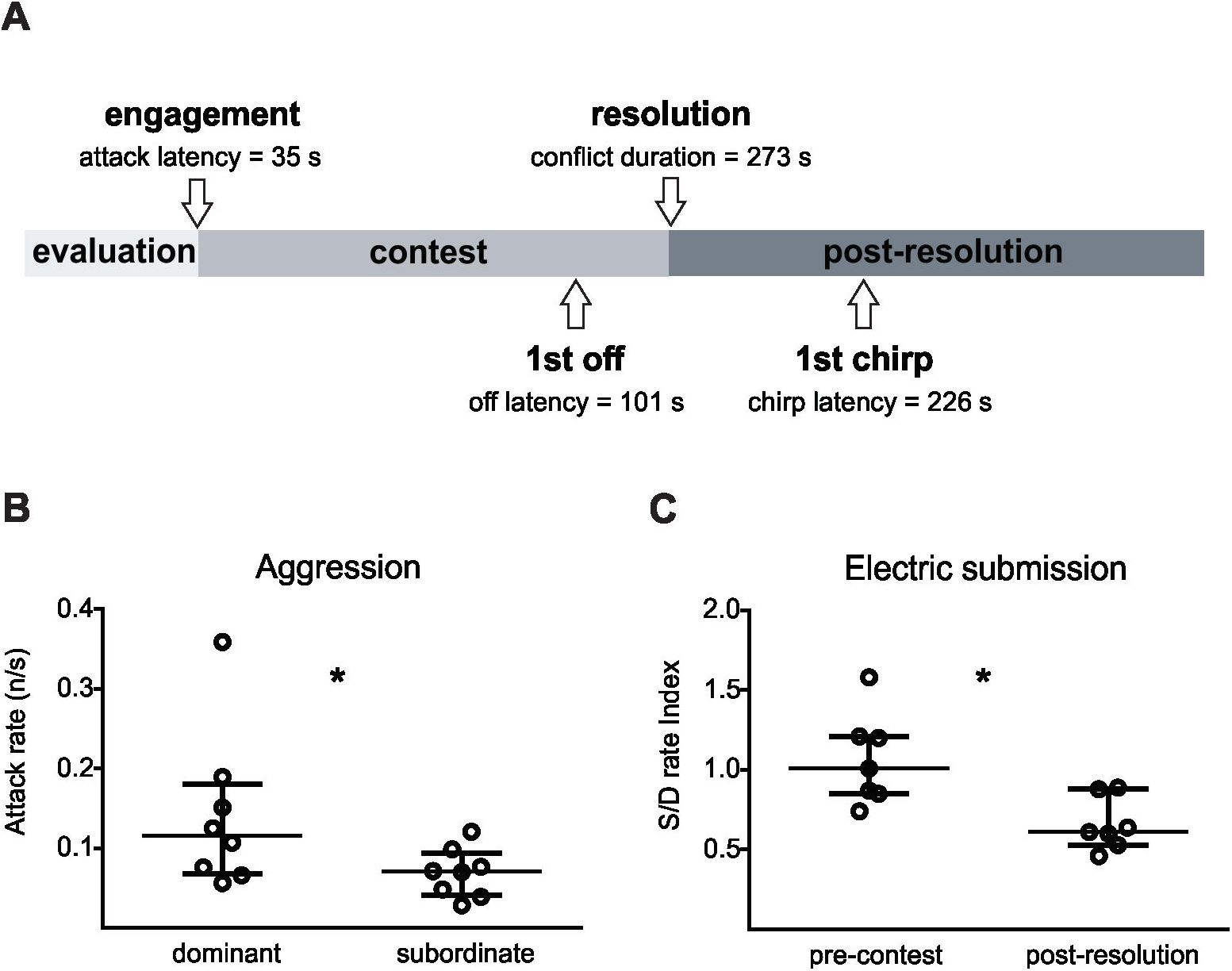
Female non-breeding aggression characterization. **A.** Female dyadic encounters displayed the three typical phases of agonistic behavior. Submission signals began during the contest phase and continued into post-resolution (n=8 control dyads). **B.** Individuals which achieved dominance showed higher aggression levels than their counterparts during conflict (attack rate Wilcoxon Matched-Pairs test, p = 0.008, n = 8). **C.** Dominant and subordinate status was expressed by post contest EOD rate. Rates were compared by the subordinate / dominant EOD rate index (S/D rate index). Index values were near 1 before contests and significantly lower after conflict resolution (pre-contest vs post-resolution S/D rate index: Wilcoxon Matched-Pairs test, p = 0.016, n = 7).

### Hormonal modulation of aggression: the analysis of rapid effects through acute treatments

To assess rapid effects of estrogens on the expression of non-breeding female territorial aggression, we acutely treated both fish of the dyad with fadrozole, an inhibitor of the aromatase enzyme. The first and foremost effect of aromatase inhibition upon dyadic interaction was a significant decline in overall aggression. As shown in Fig. 3A, 8 out of 8 control dyads engaged in conflict in less than 28 seconds; all of them reached conflict resolution and establishment of dominant/subordinate status in less than 156 seconds. Fadrozole-treated dyads showed conflict engagement in 7 of 10 dyads, of which only 5 resolved their conflict (conflict resolution: Chi square test 2×2, Fisher exact Test, p=0.035, n_FAD_ = 10, n_CTRL_ = 8). The other two dyads which engaged in conflict did not achieve resolution in a 20-minute period (Fig. 3A). Of the 5 dyads which resolved the conflict, in 3 the larger contender achieved dominance. The administration of fadrozole increased the latency to the first attack in comparison to control dyads (Mann-Whitney U test, p = 0.014, n_FAD_ = 7, n_CTRL_= 8; Fig. 3B) and in the dyads in which conflict was resolved, there was a conspicuous decrease in aggression levels. The attack rates of dominants displayed during contests were significantly lower than control dyads (Mann-Whitney U test, p = 0.019, n_FAD_ = 5, n_CTRL_ = 8; Fig. 3C), as were the attack rates of subordinate fish (Mann-Whitney U test, p = 0.006, n_FAD_ = 5, n_CTRL_ = 8; Fig. 3D). This striking overall effect upon aggression levels most probably accounts for the lower percentage of conflict resolution. However, it does not generate a significant modification in the accompanying electric social signals of submission. Subordinates of the dyads with aromatase inhibition did not differ in off rate (Mann-Whitney U test, p =0.9, n_FAD_ = 7, n_CTRL_ = 8), nor chirp rate (Mann-Whitney U test, p =0.3, n_FAD_ = 7, n_CTRL_ = 8). In addition, EOD rate rank was established in fadrozole treated dyads just as in control ones, as post-resolution S/D rate index values were lower than before the contest, reflecting a higher rate of dominants in comparison to subordinates after resolution (pre-contest vs post-resolution S/D rate index_FAD_: Wilcoxon Matched-Pairs test, p = 0.06, n = 5, data not shown). Not only did the S/D rate index decrease after resolution as in controls, but the values in themselves did not differ between fadrozole and control dyads, neither during pre-contest phase (Mann-Whitney U test, p = 0.84, n_FAD_ = 5, n_CTRL_ = 7) nor after conflict resolution (Mann-Whitney U test, p = 0.11, n_FAD_ = 5, n_CTRL_ = 7). Fadrozole treated dyads showed no difference compared to controls in locomotor activity 1 h after injection, before gate removal (Mann-Whitney U test, p = 0.28, n_FAD_ = 20, n_CTRL_ = 16, data not shown).

**Figure 3.**
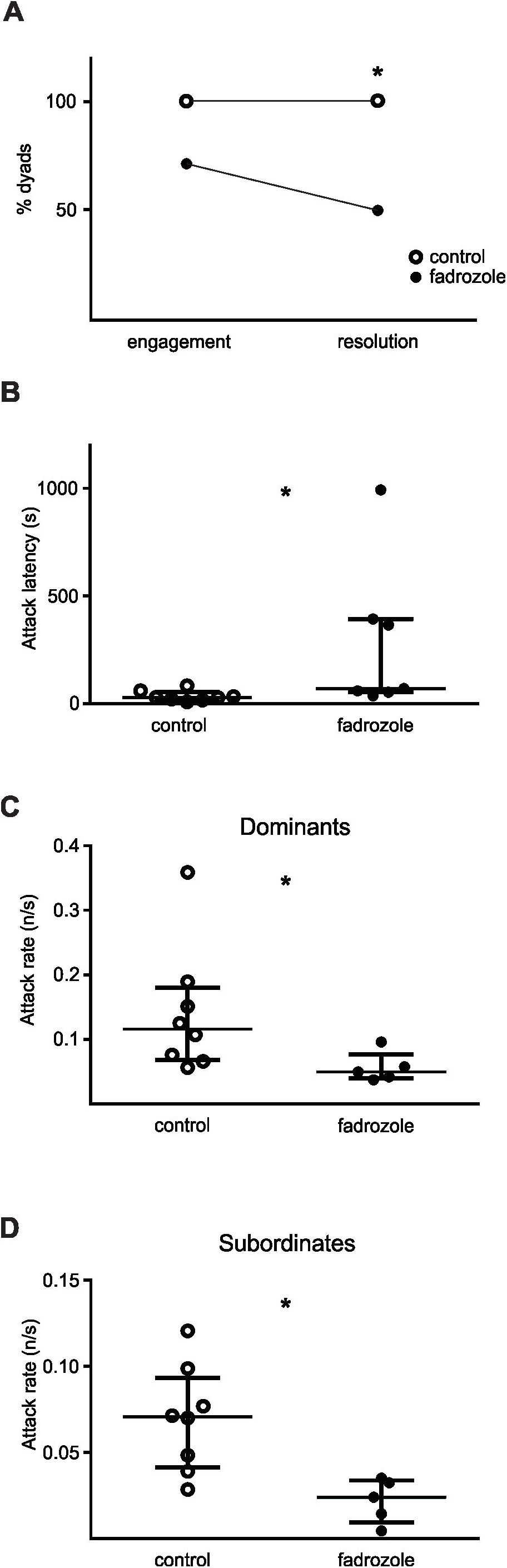
Effects of acute inhibition of aromatase on female non-breeding aggression. **A**. Fadrozole treated dyads engaged less in conflict than control dyads and only 5 dyads reached conflict resolution and established dominant/subordinate status (Chi square test 2×2, Fisher exact Test, p=0.035, n_FAD_ = 10, n_CTRL_ = 8). **B**. The latency to first attack in FAD treated dyads was significantly lower than in control dyads (Mann-Whitney U test, p = 0.014, n_FAD_= 7, n_CTRL_= 8). **C**. Individuals which achieved dominance displayed lower attack rates during contests in FAD treated dyads compared to control dyads (Mann-Whitney U test, p = 0.019, n_FAD_= 5, n_CTRL_ = 8) **D**. Subordinates showed lower attack rates in FAD treated dyads in comparison to controls (Mann-Whitney U test, p = 0.006, n_FAD_= 5, n_CTRL_ = 8).

To analyze if, in addition to estrogenic modulation, endogenous androgens have direct rapid effects upon non-breeding aggression, we treated both contenders of a group of dyads with cyproterone acetate, an androgen receptor antagonist. Acutely blocking the androgen receptor function had no effects upon overall aggression dynamics. Neither conflict engagement, latency to first attack, conflict resolution, nor aggression levels of dominant or subordinate fish showed any significant difference in comparison to control dyads (Mann-Whitney U test attack latency, p = 0.56, n_CA_ = 7, n_CTRL_ = 8; Mann-Whitney U test dominant attack rate, p = 0.67, n_CA_ = 7, n_CTRL_ = 8; Mann-Whitney U test subordinate attack rate, p = 0.09, n_CA_ = 7, n_CTRL_ = 8, data not shown). Subordinates of the dyads with cyproterone acetate did not differ in off rate emission (Mann-Whitney U test, p =0.45, n_CA_ = 7, n_CTRL_ = 8), nor chirp rate (Mann-Whitney U test, p =0.25, n_CA_ = 7, n_CTRL_ = 8). EOD rate rank was established in cyproterone acetate treated dyads as well as in control ones (pre-contest vs post-resolution S/D rate index_CA_: Wilcoxon Matched-Pairs test, p = 0.016, n = 7). Moreover, S/D rate index did not differ between cyproterone acetate and control dyads, neither during pre-contest phase (Mann-Whitney U test, p = 0.38, n_CA_ = 7, n_CTRL_ = 7) nor after conflict resolution (Mann-Whitney U test, p =, 0.38, n_CA_ = 7, n_CTRL_ = 7).

## Discussion

This is the first report on the evaluation of hormonal control of non-breeding female aggression in a teleost species. We show that a. non-breeding females of *Gymnotus omarorum* display robust aggressive territorial behavior, b. this aggression depends on rapid modulation of aromatase, revealing the importance of short-term effects of estrogens, and c. androgens show no rapid modulation upon this behavior.

Territorial aggression in *G. omarorum* has previously been reported to occur both in males and females, and be sexually monomorphic [34,35]. Nevertheless, the careful analysis of female aggression separately from male aggression is imperative to approach the hormonal modulation of this behavior. Overtly similar behavior may in fact be based on sexually distinct underlying mechanisms [49]. This is the case of the similar parenting behavior in male and female prairie voles, which are underlain by sexually different vasopressin innervation in key brain areas (reviewed in [49]). In the present study, female *G. omarorum* engaged, as expected, in highly aggressive and escalated contests during the non-breeding season, competing for space as a resource. After a short evaluation time, contests were initiated and resolved in less than 5 minutes. The larger female won most of the fights, submission was signalled by transient social electric signals and dominance was displayed by actively excluding the subordinate fish from the acquired territory while establishing an EOD rate rank, as has previously been reported [34,48,50]. Non-breeding territorial behavior in *G. omarorum* is extremely robust and maintains its features and overall dynamics independently of sex, across controls of many experimental approaches and in surgically sexed animals [33,34,36,48,50,51]. Lab results showing no sexual differences in non-breeding territorial aggression complement the data on spacing of this species in the wild, in which males and females own same-sized territories [32]. Non-breeding territorial aggression may be related to the defense of foraging patches since electrogeneration has been reported to impose high basal metabolic requirements [52]. Weakly electric fish continuously discharge EODs throughout their life and EOD amplitude is known to be strongly correlated with fish size in *G. omarorum* [53] and other electric fish [54–56]. Larger fish not only hold larger territories in the wild, regardless of sex [32], but also have a higher chance of winning a contest ([34], this study).

Non-breeding aggression in birds and mammals has been reported to be mediated by circulating precursors which are converted into active sex steroids (androgens and estrogens) within the brain (reviewed in [12]). As an exploratory step in evaluating the role of sex steroids in non-breeding female aggression in *G. omarorum*, we pharmacologically manipulated the androgenic pathway. Rapid, nongenomic actions of androgens have been reported to occur in various tissues, including the brain, mediated by androgen receptors [57–59]. Cyproterone acetate is an antagonist of androgen receptors, including those mediating fast nongenomic actions [60] and was effective in *G. omarorum* as it blocked an androgen-induced change in EOD waveform (Fig. 1). Short-term blocking of androgen receptors, however, showed no influence upon non-breeding aggression dynamics nor the establishment of dominant/subordinate status. These results suggest that if androgens are directly involved at all in sustaining aggression during the non-breeding season in females, their action may be through genomic mechanisms, and thus be evinced in a longer time frame. In male *G. omarorum*, non-breeding aggression remains unchanged under long term elimination of gonadal hormones, ruling out their role as modulators [36]. In the year-round territorial fish *Stegastes nigricans*, non-breeding circulating androgens are low, and remain so in both sexes after an aggressive encounter, although long term androgen receptor blocking decreases aggression in males but not females [17,61]. We have yet to explore if long-term direct effects of androgens, regardless of their source, occur in male and female non-breeding aggression in *G. omarorum.*

Estrogens have been put forth as key elements in models of non-breeding aggression. Pioneer studies in birds show long-lasting aromatase inhibition reduces aggression which can be recovered by estradiol treatment [24,62]. We focused on the role of this steroidal pathway in the non-breeding aggression of female *G. omarorum* using acute aromatase inhibition and showed an important role of estrogens. There was an overall decrease in motivation to display aggression, revealed both by an important delay in initiating overt aggression and a significant decrease in dyads which reached conflict resolution (Fig. 3). These results were strikingly similar to what has been reported for male *G. omarorum*, in which potential winners failed to either resolve contests or achieve dominance when acutely treated with an aromatase inhibitor [36]. Interestingly, in spite of affecting the intensity of aggressive interactions, aromatase inhibition did not affect electric signalling, which suggests that the electrogeneration system is not sensitive to rapid estrogen effects *per se*. Our results, taken together with reports of estrogenic modulation of male aggression [36] support estrogen as a key modulator of non-breeding aggression, acting through rapid mechanisms in this species. Estrogen, most probably brain derived, has been reported to have rapid effects underlying non-breeding aggression in birds and mammals [18,19,23,24,63,64]. The magnitude of these rapid effects upon behavior have been shown to depend on estrogen sensitivity i.e. higher estrogen receptor expression, in key brain regions [21]. It is interesting to focus on how female aggression can bring novel and sexually distinct mechanisms into consideration. Rapid effects of estrogens have been reported to be more pronounced in female than male brains in zebra finch [13]. Non-breeding female Siberian hamsters, which display robust aggression, have very low circulating estrogen levels which are offset by a seasonal increase in estrogen sensitivity in brain areas associated with aggressive behavior [63]. Teleost fish, which have exceptionally high aromatase activity that shows both seasonal plasticity and sexual differences (reviewed in [65]) emerge as an advantageous model for this approach.

### Concluding remarks

In this study we show for the first time in a female teleost that non-breeding aggression depends on estrogen production. Females of *Gymnotus omarorum* rely on short term estrogen synthesis to engage in territorial aggression, maintain high levels of aggression, and ultimately reach conflict resolution from which dominant/subordinate status emerges. Our results highlight the importance of fast acting estrogens in the control of non-breeding female aggression in *G. omarorum* which taken together with our reports from males of this species, as well as contributions from bird and mammal models point to conserved strategies across species. Further analysis of female non-breeding aggression may shed light on potential sexual differences in the fine tuning of social behaviors.

## Acknowledgements

We are very grateful to Cecilia Jalabert for her revision and helpful suggestions to our manuscript, and Rossana Perrone for her generous advice on behavioral processing and statistics. We would like to thank to Agencia Nacional de Investigación e Innovación (ANII), Programa de Desarrollo de las Ciencias Básicas (PEDECIBA), and Universidad de la República, Uruguay for funding.

Specific grants that funded this research: ANII_FCE_6180, ANII_FCE_136381, FCE_4272.

